# The persistence in time of distributional patterns in marine megafauna impacts zonal conservation strategies

**DOI:** 10.1101/790634

**Authors:** C. Lambert, G. Dorémus, V. Ridoux

## Abstract

The main type of zonal conservation approach corresponds to Marine Protected Areas (MPAs), which are spatially defined and generally static entities aiming at the protection of some target populations by the implementation of a management plan. For highly mobile species the relevance of an MPA over time might be hampered by temporal variations in distributions or home ranges. In the present work, we used habitat model-based predicted distributions of cetaceans and seabirds within the Bay of Biscay from 2004 to 2017 to characterise the aggregation and persistence of mobile species distributional patterns and the relevance of the existing MPA network. We explored the relationship between population abundance and spatial extent of distribution to assess the aggregation level of species distribution. We used the smallest spatial extent including 75% of the population present in the Bay of Biscay to define specific core areas of distributions, and calculated their persistence over the 14 studied years. We inspected the relevance of the MPA network with respect to aggregation and persistence. We found that aggregation and persistence are two independent features of marine megafauna distributions. Indeed, strong persistence was shown in both aggregated (bottlenose dolphins, auks) and loosely distributed species (northern gannets), while some species with aggregated distributions also showed limited year-to-year persistence in their patterns (black-legged kittiwakes). We thus have demonstrated that both aggregation and persistence have potential impact on the amount of spatio-temporal distributional variability encompassed within static MPAs. Our results exemplified the need to have access to a minimal temporal depth in the species distribution data when aiming to designate new site boundaries for the conservation of mobile species.

## 1 Introduction

Marine Protected Areas (MPAs) are spatially defined and are generally static entities aiming at the protection of some target populations through the implementation of a management plan (Kelleher, 1999). In general, MPA design aims at optimising the protection of key areas of distributions by encompassing high spatial aggregations of individuals within rather small protected areas, *i.e*. the critical habitats of target species (Hooker & Gerber, 2004). Critical habitats of a species include the habitats required for successful breeding and foraging ensuring its survival and population growth. In the case of endothermic top predators (*i.e*. seabirds and marine mammals), these critical habitats can be separated in both space and time, sometimes very distantly apart, as these species can cover thousands of kilometres per year (as for seabirds, pinnipeds or baleen whales for example; Game et al., 2009; Lewison et al., 2015).

For seabirds and pinnipeds, resting and breeding sites are well-known critical habitats, as seabird colonies and seal haul-out sites are generally well identified, and their protection is made easier by the aggregation of large amounts of individuals in restricted coastal areas (Game et al., 2009). However, the time spent within these areas is often small compared to the time spent outside, where species remain unprotected despite potentially important cumulative threats (Hooker & Gerber, 2004). Yet, both foraging habitats and access to these foraging resources are subject to a combination of major threats (acoustic and chemical pollutions, physical habitat destruction, marine debris, overfishing) and would require adequate protection. Due to the lack of knowledge about the at-sea distributions of marine top predator, especially in oceanic waters (Game et al., 2009), their protection remained poor. In the past few years, effort has been made toward extension of the coastal networks of marine protected areas (MPAs) to offshore waters in order to encompass such particular areas (*e.g*. Skov et al., 2007; Notarbartolo Di Sciara et al., 2008; Arcos et al., 2012; Garthe et al., 2012; Delavenne et al., 2017; Heinänen & Skov, 2015). This is particularly the case in the European Union where Member States are currently designating offshore sites completing the existing coastal networks of MPAs (see INPN, 2018, in France).

Although this effort of extension to the offshore top predators diversity hotspot is of crucial importance, the relevance of zonal strategies (*i.e*. establishing static MPAs) can be questioned for the conservation of highly mobile marine endothermic predators. Indeed ocean is highly dynamic in both space and time (Longhurst, 2007; Game et al., 2009), and mobile endothermic top predators are known to track the spatially and temporally varying features of interest to sustain their growth and reproduction (Ballance et al., 2006; Weimerskirch, 2007). Despite some site-fidelity linked to particular phase of their life cycle (*i.e*. reproduction, especially for seabirds or pinnipeds), habitat preferences exhibited by endothermic top predators when at-sea could be expected to vary depending on the environmental conditions experienced by species on a particular year and at a particular season (Lambert et al., 2017a, 2018). These temporally varying preferences might induce more or less important variations in distribution. For example, a breeding seabird should adjust its at-sea habitat use depending on the available environmental conditions around its colony, or odontocetes should change their distribution to match the spatial variation of their favourable habitats between years. These spatial variations in distribution might thus lead to variations in the relevance of a static MPA over years (Game et al., 2009; Lewison et al., 2015). Species with loose distribution or with strong temporal variations might more benefit from non-zonal conservation approaches, such as full national or international protection. As a result, a better understanding of the aggregation and persistence of distributional patterns of target species would ultimately help to make choice between policy instruments.

This study aimed at elucidating the effect of predator mobility on static MPA relevance in the Bay of Biscay (BoB), France, where oceanographic multi-disciplinary cruises have been conducted every spring since 2003. All seabirds and marine mammals are fully protected at the national level (against destruction, mutilation, capture, transport…) in France, but they, and their habitat, also benefit from the specific protection and conservation measures provided by various MPAs designated under diverse jurisdictional status. Seabirds are protected by Natura 2000 sites under the European Birds Directive, while marine mammals and their habitat are protected under the Natura 2000 Habitat Directive (only four species: harbour porpoise, bottlenose dolphin, grey and harbour seals). Both taxa are protected by a set of Marine Natural Parks as well. Within the Bay of Biscay, in 2018, 99 MPAs include 3 Marine Natural Parks (French Marine Natural Parks, 2019), 58 Natura 2000 sites designated under the Habitats Directive and 38 Natura 2000 sites designated under the Birds Directive (INPN, 2018). Among those Natura 2000 sites, two offshore sites of large extent have been designated in 2018 to achieve the EU Member States objectives of offshore waters protection (Delavenne et al., 2017; Journal Officiel, 2018).

We explored the implication of species mobility for zonal conservation strategies by following two main steps: (i) characterising the distributional patterns of mobile species based on two parameters, their aggregation level and their persistence; (ii) assessing the relevance of existing MPAs regarding those two parameters. We computed predictions of the distribution of eight taxa (seven seabirds, one cetacean) for each year from 2004 to 2017 in the Bay of Biscay based on habitat modelling computed from oceanographic cruise data. We identified the aggregation level of species from the relationship between population abundance and spatial extent of distribution, expecting aggregated species to have a large proportion of their population located into small areas. We defined the smallest spatial extent including 75% of the Bay of Biscay population (following a method similar to the one implemented by Nur et al., 2011) as the core area of distribution of a species, and their persistence was calculated over the 14 studied years.

Finally, we explored whether the MPA network would actually be adequate for the protection of the eight studied groups of species in respect with their core areas of distribution and their persistence, and discussed the implication of such spatially varying distributions for the conservation of mobile marine predators through static MPAs.

## 2 Material and Methods

### 2.1 Data source

This study builds from observation data obtained through the pelagic ocenographic cruises PELGAS (*PELagiques GAScogne*), conducted by IFREMER (French research institute for the exploration of the sea) onboard the research vessel *Thalassa*, which sample long transects perpendicular to the coast over the shelf every year in May/June (Figure 1a; Doray et al., 2018). Top predator observations were collected following a line transect protocol (Buckland et al., 2001) over the period 2004–2017. In-situ environmental variables were routinely collected along transects: surface and bottom temperatures, salinity, mixed layer depth and surface chlorophyll a concentration (Doray et al., 2018).

**Figure 1.**
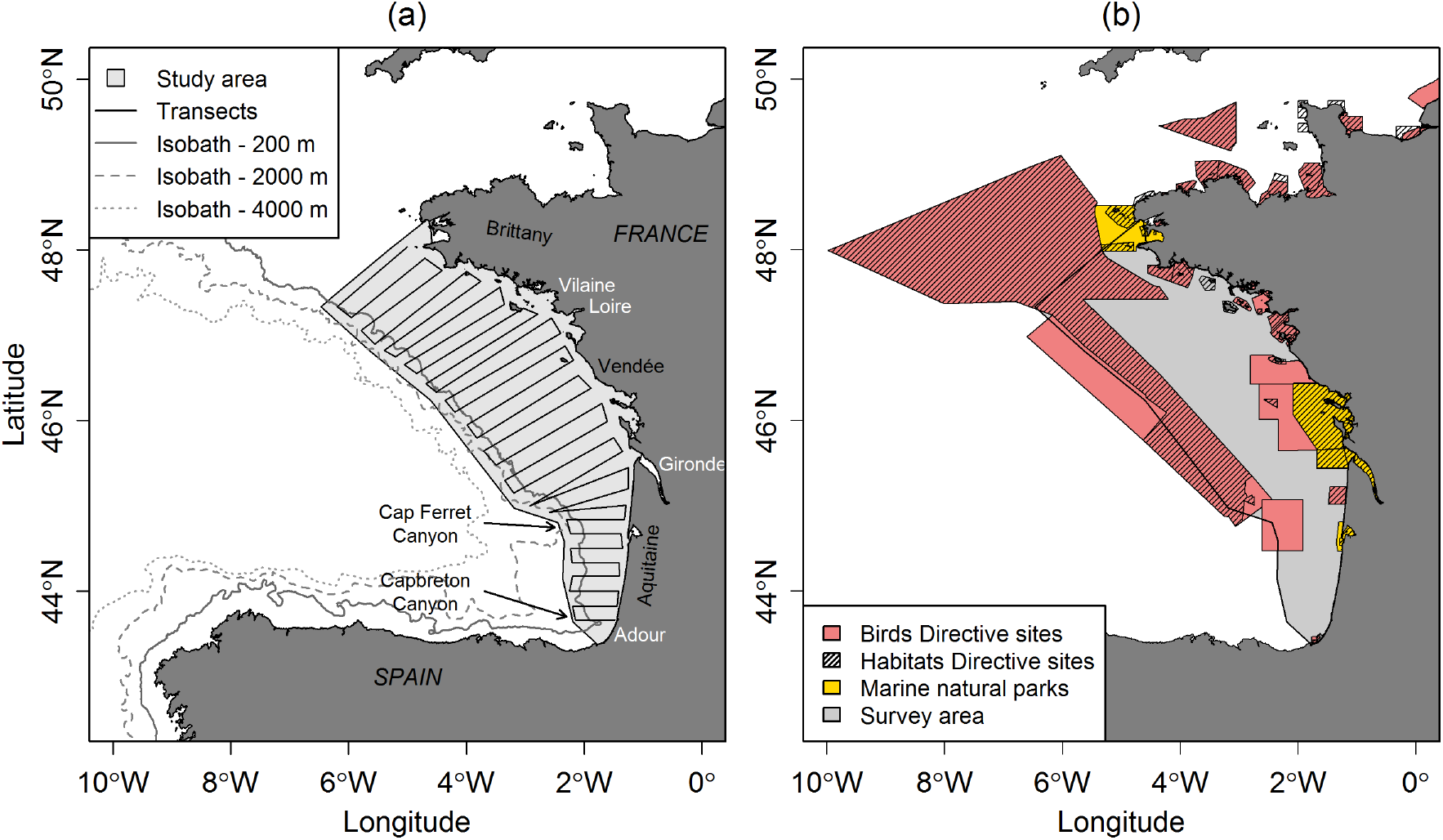
(a) Study area and theoretical sampling design of PELGAS survey. Names of four main estuaries in white, names of other geographical localities and main canyons in black; (b) French Marine Protected Areas (MPAs) in the Bay of Biscay and English Channel. This study considered only sites overlapping the survey area. Birds Directive sites were assessed only for seabirds, Habitats Directive sites only for cetaceans.

Our analysis builds from the habitat models produced by and described in (Lambert et al., 2018). The habitat modelling procedure mostly used environmental variables collected *in-situ* during the oceanographic cruises, but also some environmental variables derived from remote-sensing sources and from bathymetric grid of the ocean (Appendix A). The procedure takes into account the variability of habitat preferences across years by selecting between a global model (considering the relationship with environmental variables similar over years) and an interaction model (integrating the interaction between variables and years, allowing the relationship to change between years). The habitat modelling procedure is detailed in the Appendix A, along with maps of predicted densities in individual per square kilometres for each studied years (2004–2017) for the eight taxa.

Here, we discarded the common dolphin from the set of studied species from (Lambert et al., 2018) as habitat modelling failed to predict correctly their distributions, but considered three more taxa, whose distribution were predicted following the same procedure. During at-sea data collection, some individuals are impossible to tell apart between closely related species exhibiting close morphology and behaviour. As a result, we focused on a set of individual species and taxa composed of several closely related species: bottlenose dolphin (*Tursiops truncatus*); northern fulmar (*Fulmarus glacialis*); small-sized shearwaters (Manx *Puffinus puffinus* and Balearic *P. mauretanicus* shearwaters); storm petrels (European *Hydrobates pelagicus*, Leach’s *H. leucorhous* and band-rumped *Hydrobates castro* storm-petrels); northern gannets (*Morus bassanus*); great skua (*Catharacta skua*); auks (common guillemot *Uria aalge* and razorbill *Alca torda*); black-legged kittiwake (*Rissa tridactyla*).

### 2.2 Distribution patterns

#### 2.2.1 Aggregation level

We first transformed density maps from habitat models to abundance maps by multiplying predicted density by cell surface. The abundance maps were then transformed into proportion maps, *i.e*. the abundance of each cell was related to the total abundance predicted within the study area (sum all cells within the PELGAS stratum). For each species and each year, all cells were sorted by decreasing predicted proportions and the cumulative sum was computed. We explored the aggregation level of each species distribution by plotting the cumulative sum of abundance proportion against the corresponding cumulative sum of surface for each species and year. Aggregated species were identified as species with high proportions of population concentrated into small surface *versus* larger surface for broadly distributed species (relationship closer to linearity).

#### 2.2.2 Core areas of distribution and their persistence

The smallest spatial extent including 75% of the population identified the core areas of distribution of studied taxa. Based on the cumulative sum of abundance proportion, the set of cells containing 75% of the population was assigned the value of “1”, all remaining cells were assigned “0”.

The persistence of core areas was calculated as the number of years each cell belonged to the core area (category 1). Habitat models being built on *in-situ* variables, some cells have no prediction for years during which they were not sampled. To take into account this variation, the number of years a cell belonged to the core area was divided by the number of years each cell was sampled. The persistence was thus expressed as the proportion of sampled years a cell was included in the core area of distribution.

### 2.3 MPA relevance within the Bay of Biscay

The proportions of core areas of distributions actually falling within MPAs for each studied year was quantified to assess the relevance of MPAs within the Bay of Biscay: we considered all cells of the core area whose centre was inside an MPA as included in that MPA. We quantified as well the proportions of persistent cells (*i.e*. belonging to the core area at least 50% of surveyed years) whose centres fall within MPAs.

We assessed Bird Directive sites for seabirds, and Habitats Directive sites for bottlenose dolphin (Figure 1b). The Bird Directive sites target the protection of bird species, while the Habitats Directive sites aim at protecting, among other species, the bottlenose dolphin habitat. We only considered sites overlapping with the study area.

## 3 Results

### 3.1 Habitat modelling

The interaction model was selected for most species (all but bottlenose dolphins and storm petrels; Appendix A), indicating their relationship to their habitat might vary to some extent across years. Selected models resulted in reasonably good explain deviances (from 21.7% for storm petrels to 58.9% for black-legged kittiwakes, with an average of 40.5%) and fitted well the observed distribution of species across years.

The seven studied taxa exhibited different predicted distribution patterns (Figure 2; see Appendix A for yearly predicted distributions). Bottlenose dolphins exhibited the less variable and most aggregated spatial distribution, with a very clear preference for the shelf edge (Appendix A3). Kittiwakes also exhibited an aggregated distribution, with highest predicted densities along the coast of northern BoB (Appendix A4). Auks were the third most aggregated taxon, occurring mostly along the coast during the fourteen years, especially within river plumes (Appendix A5). Storm petrels were predicted over the whole northern BoB shelf and offshore Basque country (Appendix A6). The northern fulmars were mostly predicted over the slope and outer shelf of the BoB, avoiding coastal areas, during the fourteen years (Appendix A7). The distribution of small-sized shearwaters varied more between years, but they remained mostly predicted in inner and central shelf areas of the northern BoB (Appendix A8). Northern gannets were distributed over the whole BoB, with higher densities in the northern part (Appendix A9).

**Figure 2.**
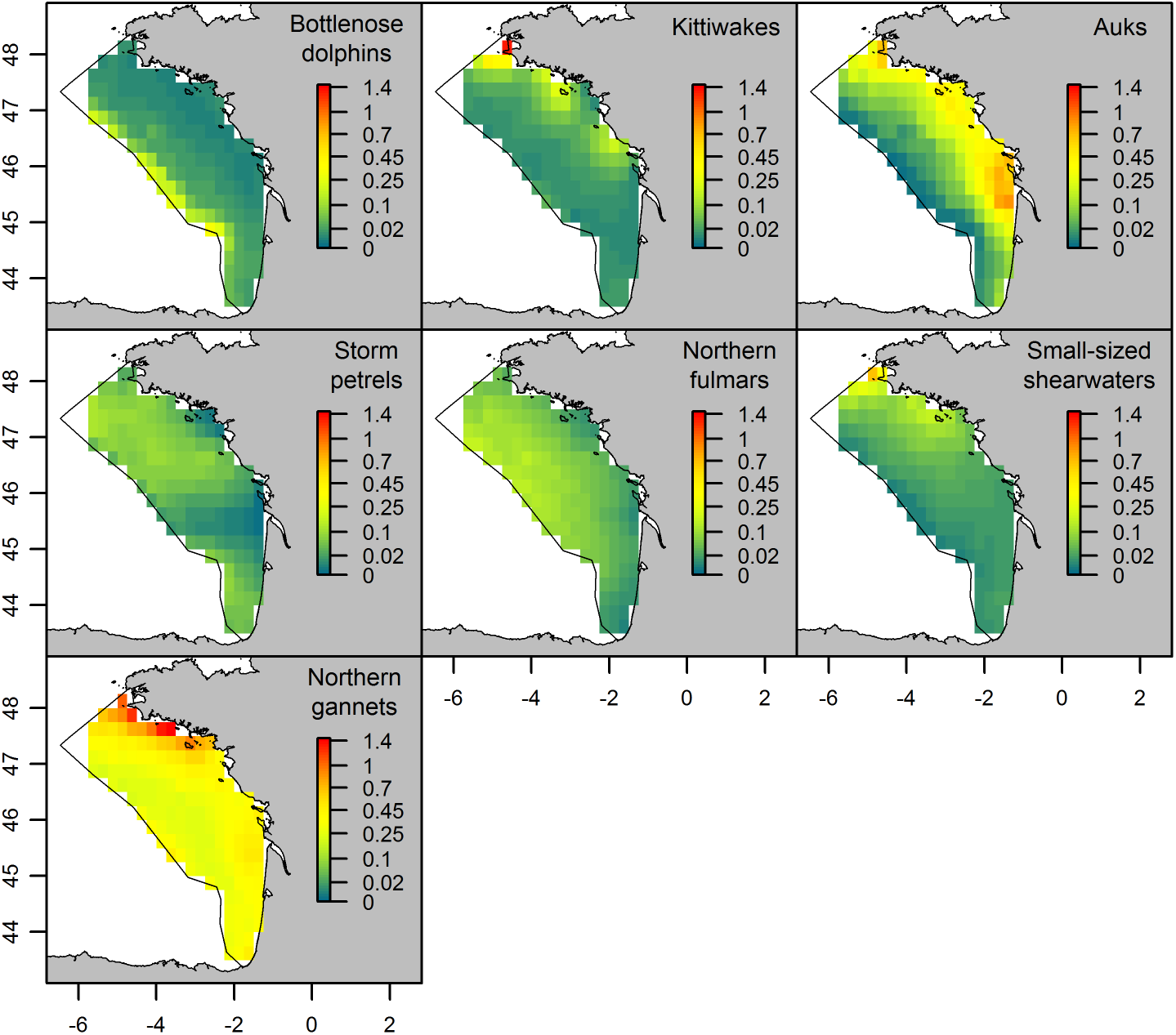
Predicted densities (individual/km^2^) averaged over the fourteen studied years for the seven studied taxa.

### 3.2 Aggregation levels

We expected species with an aggregated distribution to have high proportions of population in small surface compared to species with a loose distribution.

Bottlenose dolphins, black-legged kittiwakes and auks presented the highest aggregation levels among the studied species, their core areas showing the smallest spatial extent (Figure 3). In average, 75% of the population was encompassed within 20, 23 and 25% of the study area, respectively. The curves rapidly reached this value, then the proportion of population levelled off with the increase of stratum surface proportion. The aggregation level varied somehow across years for bottlenose dolphins and auks, but the overall relationship remained the same throughout the studied years (Figure 3). Bottlenose dolphins core areas were restricted to the shelf edge during all the studied years (Appendix B1), while auks were mostly aggregated over river plumes from the Vilaine to Gironde estuaries, with some years core areas occurring within the Adour river plume (2004–2009; Appendix B3; see Figure 1 for location of these estuaries). However, the black-legged kittiwake distribution showed different pattern of aggregation during three of the studied years, being highly aggregated in a few cells in some years (2017), but broadly distributed in two others (relationships tending toward linearity, 2005–2006; Figure 3). Overall, black-legged kittiwakes were mainly aggregated in southern Brittany, with an extension down to the Gironde estuary during some years (Appendix B2).

**Figure 3.**
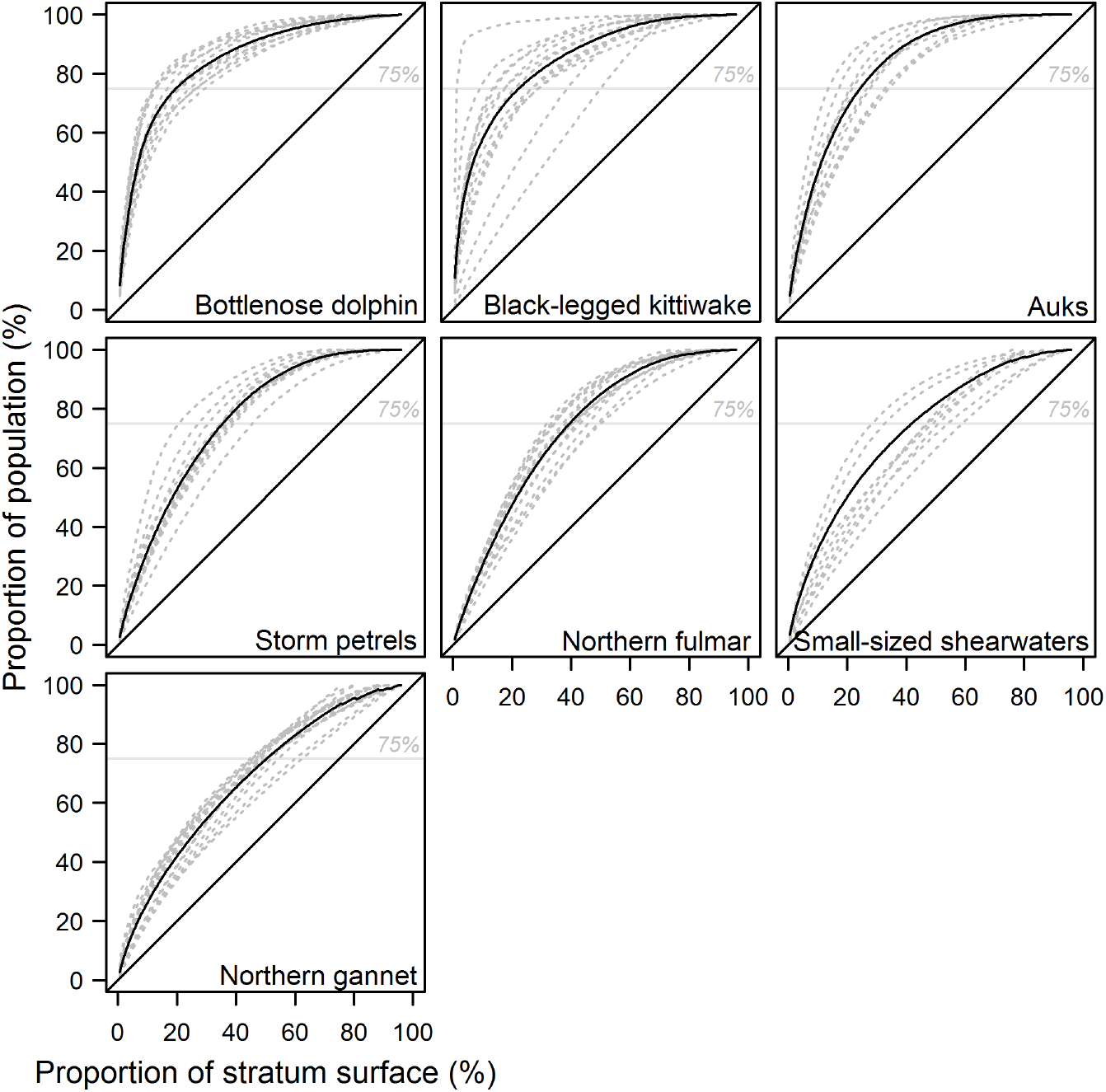
Relationships between the proportions of population covered and the corresponding proportions of stratum surface for the seven studied taxa. Annual relationships are dotted grey lines, the averages over the fourteen years are plain black lines. The 75% of population threshold used to determine core areas is shown in grey.

An intermediate aggregation level in distribution was observed for storm petrels, northern fulmars and small-sized shearwaters, with, in average, 75% of the population included in 36, 39 and 42% of the study area, respectively (Figure 3). A similar pattern was observed for all studied years, showing only limited variations, for storm petrels and northern fulmars. Storm petrels had a main core area located in the northern part of the BoB, whose extent varied somehow across years, and a secondary one over the shelf edge of the southern BoB during some years (Appendix B4). Northern fulmar core areas were consistently located over the outer shelf of the northern part of the study area (Appendix B5). The small-sized shearwaters aggregation level increased throughout the studied years (Figure 3) due to a contraction of their core areas in the north of the study area (Appendix B6). The spatial extent covered by 75% of the population shifted from 50–60% of the study area (broad distribution, with a relationship tending toward linearity) in 2004–2010 to 21% in 2017.

Northern gannets exhibited broad distribution with reduced aggregation level (Figure 3; Appendix B7). Their relationships between population and surface was almost linear, with very few variations across years. In average, 75% of the population occupied 45–60% of the study area.

### 3.3 Persistence of core areas

Bottlenose dolphins exhibited the largest spatial consistency across years, and their core area of distribution was strongly persistent (Figure 4a): the bottlenose dolphin core area of distribution (representing only 21% of the study area; Figure 4b) was located over the shelf edge 100% of surveyed years, and the vast majority of the BoB was never encompassed within the species core areas. Kittiwakes had the lowest core areas persistence due to the spatial variation of its core area across years (Figure 4a). Kittiwakes were nevertheless located off Brittany and along the Vendée coast during more than 50% of the studied years, which represented 21% of the study area (Figure 4b). The extreme north of the BoB had a persistence larger than 80% of years, but those cells were sampled during less than 10 of the studied years. Auks’ core areas had strong persistence, with estuaries being included in a core areas more than 50% of surveyed years (Figure 4a), resulting in a persistent area (*i.e*. more than 50% of surveyed years) representing 29% of the study area (Figure 4b). The rest of the shelf was never used by auks. The storm petrels’ core area located in the northern BoB was persistent across years (>80% of surveyed years; Figure 4a). The second core area, located in the southern BoB, was a bit less persistent (about 50% of surveyed years). The persistence of northern fulmar core areas was high, with a large area over the outer shelf being included in core areas more than 90% of years the cells were surveyed (Figure 4a). The same occurred for small-sized shearwaters, for which a large amount of cells was persistent more than 80% of surveyed years, from the southern Brittany to the Gironde estuary (Figure 4a). For those three species, 45, 45, 48% of the survey area was persistently included in core areas (*i.e*. more than 50% of surveyed years; Figure 4b). The northern gannets were widely distributed over the BoB across all the years, and all cells were included in a core area at least during one year (Figure 4a). The most persistent areas were located off Brittany and off the Gironde estuary. 65% of the Bay of Biscay belonged to a core area for at least 50% of surveyed years (Figure 4b).

**Figure 4.**
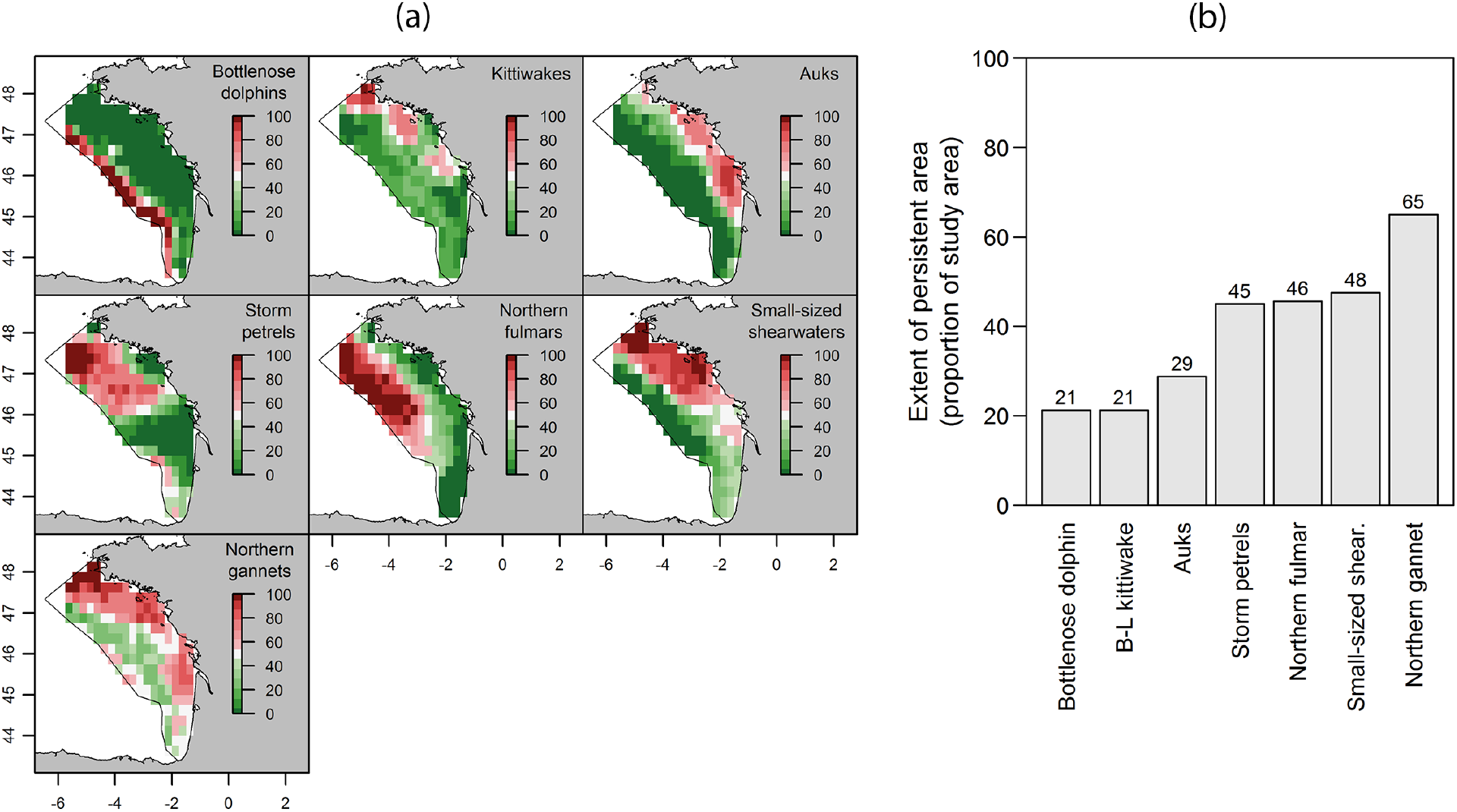
(a) Persistence of core areas of distributions by species. The persistence is expressed as the proportion of surveyed year a cell was included in species’ core area of distribution (in percent). (b) Spatial extent of persistent area by species, expressed as the proportions of the study area included in core areas more than 50% of the studied years, in percent.

### 3.4 MPA relevance within the Bay of Biscay

In the BoB, the Bird Directive sites are currently covering 68% of the stratum cells, Habitats Directive sites 58%. The above-identified core areas of distribution covered variable proportions of the study area, depending on species but also depending on years. The aggregation levels of species and the location of core areas led to varying amount of core areas being actually included within MPAs.

Thanks to their aggregated and persistent distribution over years, the proportion of bottlenose dolphin core areas within MPAs did not vary much, but was quite high thanks to the new offshore Habitats Directive site covering the shelf edge (42–67%; Figure 5). The proportion of core areas of black-legged kittiwakes in MPAs were highly variable across years, from 21% to 80% (100% in 2017 when the core area was made of only 1 cell; Figure 5), as a result of their core areas being poorly persistent. Auks were among species with aggregated distribution persistent over time, resulting in fairly high proportion of core areas in MPAs, from 35 to 68% (Figure 5). Storm petrels, northern fulmars, small-sized shearwaters were more widespread, their larger core areas being well persistent. As a result the proportion of their core areas in MPAs were medium, and showed low variation across years (24–35% for storm petrels, 33–51% for northern fulmars, 19–37% for small-sized shearwaters; Figure 5). Northern gannets were widespread, with large core areas, but these showed some variations in distribution across years, leading to variable proportions of core areas covered by MPAs, from 25 to 59% in MPAs (Figure 5).

**Figure 5.**
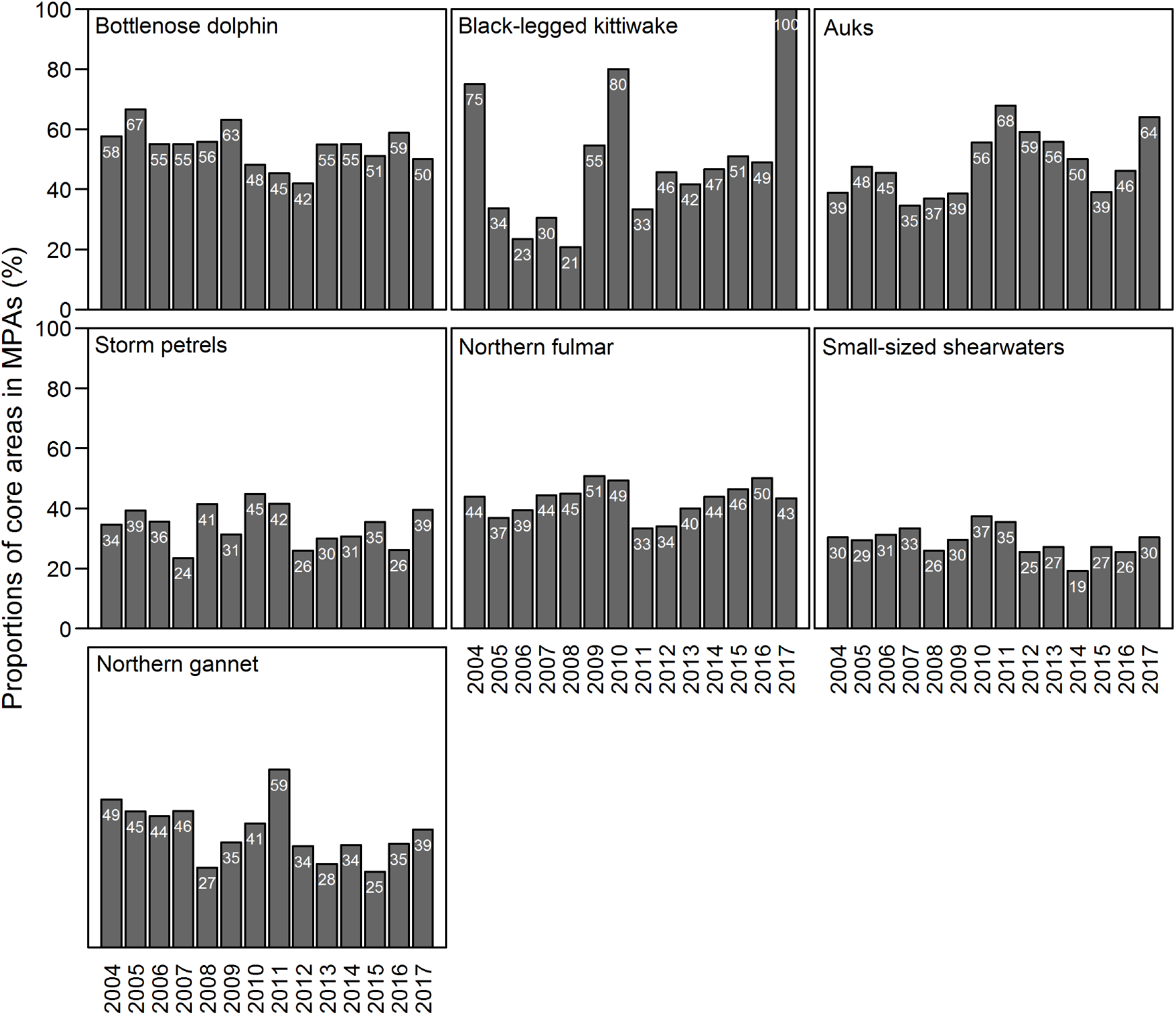
Proportions of core areas encompassed within existing MPAs (Marine Natural Parks and Bird Directive sites for seabirds; Marine Natural Park and Habitat Directive sites for bottlenose dolphins) along the fourteen years for the eight studied groups of species, in percent. The proportion is indicated in each bar.

The proportions of persistent area (*i.e*. area included in core areas more than 50% of the studied years) included in MPAs varied across species (Figure 6). Thanks to the strong persistence and aggregation of their core area over the shelf edge, 59% of the bottlenose dolphin persistent area was included in MPA (the largest proportions among studied species; Figure 6). Black-legged kittiwakes, auks, northern fulmars, small-sized shearwaters and northern gannets showed similar medium proportions of persistent area in MPAs (34–49%, Figure 6). Storm petrels persistent areas were the least covered by MPAs, with only 32% (Figure 6).

**Figure 6.**
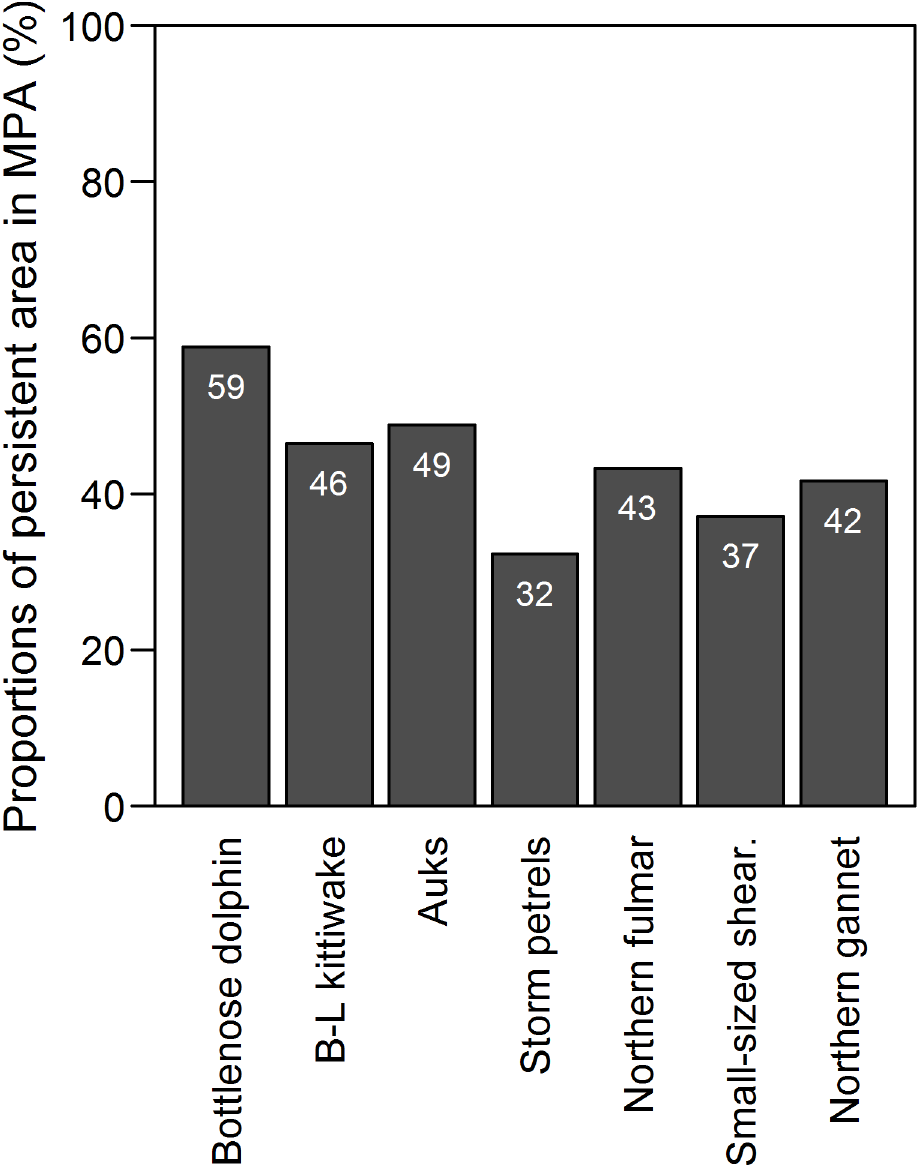
Proportions of persistent areas (cells included in core areas more than 50% of the studied years) encompassed within existing MPAs (Marine Natural Parks and Bird Directive sites for seabirds; Marine Natural Parks and Habitat Directive sites for bottlenose dolphins) for the eight studied species, in percent. The proportion is indicated in each bar.

## 4 Discussion

The Bay of Biscay is a rather small area compared to some offshore MPAs implemented worldwide, such as the Pelagos Marine Sanctuary or the Coral Sea Commonwealth Marine Reserve, but the surveys conducted annually since 2004 were a unique opportunity to investigate some of the main limitations of zonal conservation strategies for mobile species (Game et al., 2009; Wilson, 2016). Those limitations are linked to the mobility of animals, in terms of movements and relationships to habitat, but also to the variability of pelagic habitats, which are hard to characterise and highly dynamic in space and time. These combined factors lead to the conclusion that for such species, a relevant protection would necessitate larger areas as movement rates increase (Hooker & Gerber, 2004; Lewison et al., 2015). However, marine predators represent a wide range of species with various distributional patterns, and many species are known to target discrete and predictable oceanographic features (Ballance et al., 2006; Weimerskirch, 2007). Such aggregated species might well benefit from zonal conservation approaches (Oppel et al., 2018). In addition, some species might be more vulnerable within a small proportion of their range. All these elements make possible to meet conservation objectives by focusing on a few critical areas (Game et al., 2009).

Here, we aimed at investigating the effect of temporal variability in distribution for a set of marine top predators on the potential protection by static MPAs. First, we were successful in characterising the habitats available within our study region thanks to the use of a PCA based on *in-situ* environmental conditions monitored simultaneously to the megafauna survey (Lambert et al., 2018). The quality of models was reasonable to fairly good regarding the standard of habitat modelling for these organisms (good deviances and good predictions–sightings adequacy; see for comparison: Vilchis et al., 2006; Becker et al., 2014; Breen et al., 2017; Lambert et al., 2017a). The habitat modelling highlighted a range of habitat strategies based on the specificity and inter-annual stability of species preferences. Species exhibiting narrower habitat preferences also exhibited stronger stability in their preferences among years (*e.g*. bottlenose dolphins and auks) while the species with wider habitat preferences exhibited higher variability among years (*e.g*. northern gannets; see Lambert et al., 2018). This variability of preferences might either originate in species being flexible in their preferences or in the actual seasonal timing of the pelagic ecosystem varying in some extent across years, ultimately driving some differences in relationships of species to their habitat. Yet, despite this possible variation in seasonal timing, the oceanographic cruise always occur during the reproductive period of studied species. As such, we are confident that the conclusion presented here regarding the impact of the persistence of distributional pattern on zonal conservation are reliable for adjusting conservation measures during this highly critical period that is the breeding period.

Of course the results presented would benefit from a similar study to be implemented over the rest of the year. Indeed, previous study highlighted seasonal variations of habitat preferences and distribution of cetaceans and seabirds in the Bay of Biscay (Lambert et al., 2017a; Laran et al., 2017; Pettex et al., 2017), showing limited seasonal differences in distribution for bottlenose dolphins and northern fulmars, but large differences in distribution for black-legged kittwakes, auks and northern gannets between summer and winter. The small-sized shearwaters are completely absent from the study region during winter. An extension of the present study to other seasons would mostly benefit to black-legged kittiwakes, auks and northern gannets, as their difference in distribution is in part due to the individuals present in the study area during winter being of different population than during summer. Unfortunately however, we still lack dataset with sufficient temporal depth to replicate the present study during other seasons.

Given the observed range of habitat strategies exhibited by taxa studied during their breeding period, we found various levels of temporal variability in aggregation and location of core areas according to the species. The relationship between the proportion of population and surface clearly identified several species with aggregated distributions on small areas, such as bottlenose dolphins, kittiwakes and auks (75% of the population was concentrated over 22% of the area, in average), and other species with broader distributions, such as northern gannet (75% of the population was spread over 50% of the area, in average). As such, we confirm that for zonal conservation to be effective for a target species, the latter needs to have an aggregated distribution (Oppel et al., 2018), but these areas of higher density must also be persistent in time. Our results showed varying patterns depending on species,highlighting that aggregation and persistence do not always covary: bottlenose dolphins and auks exhibited aggregated distribution with strong persistence over the decade; storm petrels, northern fulmars and northern gannets were widespread species with medium to high persistence but black-legged kittiwakes were an aggregated species with low persistence.

Theoretically, species with more persistent distributions should be the easiest to protect with zonal conservation strategy, and the more the distribution is aggregated, the smaller the required protected area would have to be. In our case, it would be possible to design MPAs based on the persistent distributional patterns for bottlenose dolphins, auks, storm petrels, northern fulmars, small-sized shearwaters and northern gannets. The resulting MPA would be fairly small for bottlenose dolphin and auks, thanks to their aggregated distribution, but would be larger for storm petrels, northern fulmars and small-sized shearwaters (50% of the study area). In case of aggregated species with lower persistence (black-legged kittiwakes) and species loosely distributed with important persistence area (northern gannets) in contrast, the establishment of a zonal conservation would necessitate a large MPA, to encompass all the observed temporal variability in core area distributions in one case, to encompass the whole persistent area in the other case. Those species might benefit more from non-zonal conservation approaches, such as national or international regulation of incidental mortalities linked to fisheries bycatches, or extraction of foraging resources at a larger scale for example. In the Bay of Biscay in particular, all species benefit from a generic national and european-level protection from direct destruction, in addition to the particular conservation measures implemented in MPAs.

Here, our goal was not to propose new sites, since many MPAs already exists which currently cover 68% of the study area for the Bird Directive sites, 52% for the Habitats Directive sites. The investigation of the overlap between species core areas and these MPAs showed that bottlenose dolphins and auks, the two most aggregated taxa with strong persistence, had the highest coverage by MPA with reduced temporal variability. This was achieved through the important coastal network of MPA for auks. The boundaries for Habitats Directive and Bird Directive sites were historically proposed mostly based on expert’s knowledge of coastal distributions, with poor information on their temporal variability, and *a fortiori* on the target species at-sea distributions leading to a succession of small and large sites along the BoB coast, ensuring a good coverage of the auks distribution. The important coverage of bottlenose dolphin distribution (59% of its core area) was largely ensured by the new offshore Habitats Directive site (see Figure 1) that has recently been designated, along with an equivalent Bird Directive site, based on dedicated large-scale surveys (SAMM surveys; Lambert et al., 2017a; Laran et al., 2017; Pettex et al., 2017) within French waters to compensate for the previous absence of any protected sites within offshore waters (Delavenne et al., 2017).

Our results demonstrate the interest of these new sites for the bottlenose dolphin, as they included most of its range, but also for northern fulmars and storm petrels. Those two latter taxa were more broadly distributed than the bottlenose dolphin, with larger core areas strongly persistent over the outer shelf. Prior to the designation of the offshore sites, they were as poorly covered by the coastal network of MPAs as the bottlenose dolphin (Lambert et al., 2017b), but here, we demonstrated that the offshore sites contain an important proportion of their persistent core areas (32% of storm petrels, 43% of northern fulmars). The BoB slope has recently been identified as an area with important densities of marine species whose distributional range up to now poorly overlapped with any MPAs (Klein et al., 2015). Among marine species, mammals are the species group with the lowest proportions of species range overlapping with MPAs. Our results demonstrated that the designation of the two new large offshore sites was a crucial advance toward the protection of species with offshore distribution (both mammals and seabirds), but remains to be confirmed by the establishment of an efficient management plan, a work in progress at present.

Despite these positive points, we showed that fairly large proportions of the core persistent areas (more than half) fell outside MPAs in our study area for all species but bottlenose dolphins. Yet, the BoB belongs to the ocean’s most impacted areas by cumulative human impacts (Halpern et al., 2008, 2015). We can thus wonder whether these medium to low levels of protection represent a brake to the effectiveness of conservation strategies implemented within the BoB. In his recent editorial, Wilson (2016) argue that the lag between the identification and the designation of MPAs would inevitably lead to a drop of densities within MPAs, due to the dynamic drivers of species distributions and to their mobility inducing temporally varying distributional patterns, as shown here. However, the protection of half of a species core area is surely better than providing no protection at all, especially if the protected areas cover core area with higher threats or species vulnerability (Game et al., 2009): several case studies have shown that protecting critical habitats or reducing area-specific threats can strongly reduce overall mortality rates in spite of the mitigation action taking part on a small part of the species ranges (*e.g*. Hyrenbach et al., 2006; Alpine & Hobday, 2007). Therefore, despite the intermediate to limited proportions of core and persistent areas of species distributions within MPAs in the BoB, the target species should theoretically benefit from the implemented zonal conservation strategies. This is particularly true for offshore distributed species that had very low level of zonal protection before the establishment of the two offshore sites (Lambert et al., 2017b). Obviously, the assessment of the actual efficiency of those boundaries would be completely dependent on the relevance and efficiency of the management plans to be defined and implemented within each single MPAs (Edgar et al., 2014) and remains to be addressed at the BoB scale.

## 5 Conclusion

Our results showed varying levels of temporal persistence in distributional patterns according to predator species combined with various levels of aggregation in distribution. The important result here was that these two factors did not necessarily covary, since strong persistence was shown in both aggregated and loosely distributed species, while some species with aggregated distributions also showed limited year-to-year persistence in their patterns. As a consequence, we have demonstrated that these two factors have potential impact on the amount of spatio-temporal distributional variability encompassed within static MPAs implemented over the study area. Our results exemplified the need to have access to a minimal temporal depth in the species distribution data when aiming at designating new site boundaries for the conservation of mobile species, as this would be the only way to minimize the bias linked to the species and environment mobility (as discussed by Game et al., 2009; Wilson, 2016).

## Data accessibility

Data from the PELGAS surveys used for this analysis are freely accessible on the Sea scientific open data edition (SEANOE) repository at http://doi.org/10.17882/53389 (Doray et al., 2018).

## Acknowledgements

CL was funded by the French ministry in charge of research (*Ministère de l’Enseigmenent Supérieur et de la Recherche*, MESR) during her PhD. We are indebted to crew members on-board the R/V Thalassa, to all observers who participated in the surveys as well as to the French Office of Biodiversity (*Office Français de la Biodiversité*, OFB) and French ministry of environment (*Ministère de la Transition Écologique et Solidaire*) who funded the megafauna observers. Paul Bourriau and Martin Huret are thanked for taking care of the CTD casts and maintaining the hydrological sensors, Matthieu Doray for his coordination of the survey. Version 3 of this preprint has been peer-reviewed and recommended by Peer Community In Ecology (https://doi.org/10.24072/pci.ecology.100048).

## Appendix

The Appendices A and B are available online at https://doi.org/10.1101/790634.

