## Appendix A for "The persistence in time of distributional patterns in marine megafauna impacts zonal conservation strategies"

### Habitat modelling

The study presented in this paper builds from habitat modelling presented in Lambert *et al.* (2018), extended with four supplementary years in data: Lambert *et al.* 2018 worked on data covering the 2004-2013 period, but in the present paper we worked on data covering the 2004-2017 period.

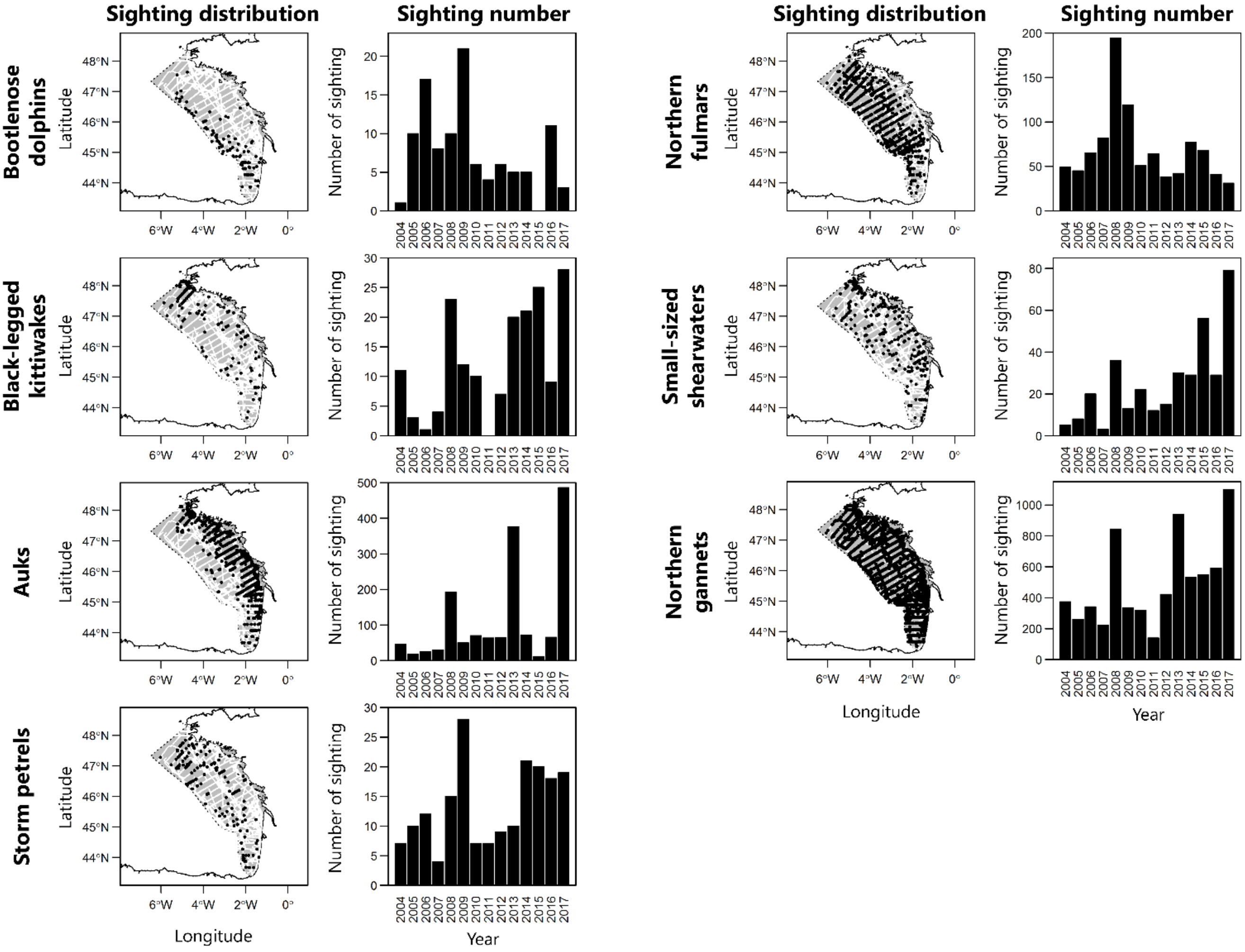


**Figure 1. Distribution of sightings in the study area and the number of sightings per year for the studied taxa.** Individual sightings in black, survey effort in white.

The habitat modelling was done over a 0.25° grid aggregating both the environmental variables and the survey effort conducted in good conditions (Beaufort sea state < 4 and medium to excellent observation conditions). The modelling used a set of nine environmental variables, of different origins. Sea surface temperature (SST), sea bottom temperature, mixed layer depth and sea surface salinity were measured directly *in-situ* during the oceanographic cruise. SST gradient was computed from SST as the largest difference between each cell and its neighbours. The surface concentration in chlorophyll *a* and net primary productivity were extracted from monthly MODIS composites for each year (http://oceancolor.gsfc.nasa.gov for chlorophyll a and http://www.science.oregonstate.edu/ocean.productivity for net primary productivity). We used the bathymetry and slope from the GEBCO 08 database.

At first, a Principal Component Analysis (PCA) was computed over these variables to explore the structuring of the environmental conditions over the study area.

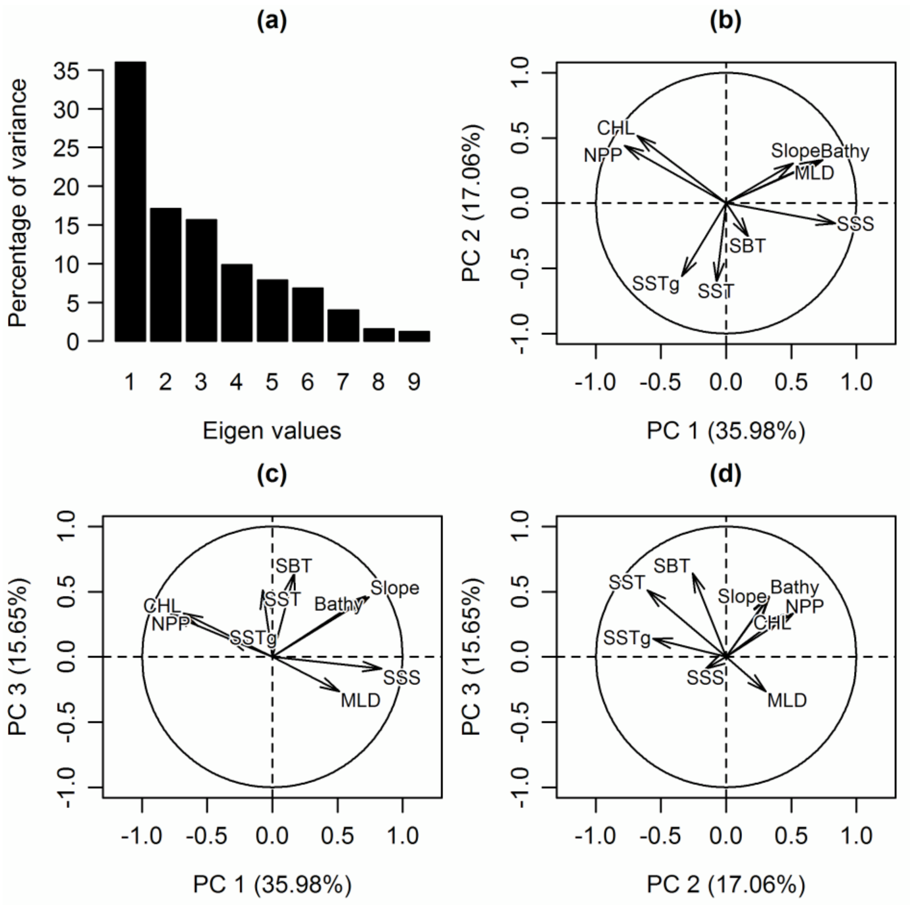


**Figure 2. Results of the Principal Component Analysis (PCA)**: (a) percentage of variance explained by each of the nine eigen values; (b) PCA correlation circle for the first and second Principal Components (PCs); (c) PCA correlation circle for the first and third PCs; (d) PCA correlation circle for the second and third PCs. SST: Sea Surface Temperature; SSTg: SST gradients; SBT: Sea Bottom Temperature; SSS: Sea Surface Salinity; MLD: Mixed Layer Depth; CHL: surface chlorophyll a concentration; NPP: Net Primary Production.

Second, we used the PCA dimensions as covariates for modelling the habitat. This was done with Generalized Additive Models (GAMs; Wood 2006), relating the number of sighted individuals and the covariates, considering the sampled area per cell as an offset and using a log-link function of the tweedie family (Foster and Bravington 2013). The covariates were the three PCA dimensions, but we also considered the distance to the closest colony for seabird species. The model selection procedure takes into account the variability of habitat preferences across years by selecting between a global model (considering the relationship with environmental variables similar over years) and an interaction model (integrating the interaction between variables and years, allowing the relationship to change between years). The model with the lowest AIC and predictions fitting best the sighting data was retained as the best model (Table 1). When the AIC difference between the two models was negligible, we parsimoniously selected the simplest model as best model (the case of storm petrels, Table 1). We predicted from selected models the distribution of each taxa for each years (2004 to 2017) over the study area, in density in individuals per km² (Figure 3-10). The whole habitat modelling procedure was computed in R 3.4.3 (R Core Team, 2017).

**Table 1. Generalized Additive Model results for the studied taxa.** The explained deviances (in %) and the significance levels for the four covariates are given, as well as the model AICs for the global and the interaction model. PC: Principal Component. ***: p-value ≤ 0.001; **: p-value ≤ 0.01; *: p-value ≤ 0.05. Selected model for each taxon is indicated by grey-shaded cell and bold font.

| | | Bottlenose dolphins | Black-legged kittiwakes | Auks | Storm petrels | Northern fulmars | Small-sized shearwaters | Northern gannets |
|---|---|---|---|---|---|---|---|---|
| Global model | Explained deviance (%) | **34.8** | 46.0 | 35.7 | **21.7** | 20.9 | 32.9 | 23.4 |
| | PC 1 | *** | * | *** | ** | *** | | *** |
| | PC 2 | * | | ** | * | | ** | |
| | PC | *** | *** | *** | ** | *** | ** | *** |
| | Distance to closest colony | - | *** | *** | *** | *** | *** | *** |
| | AIC | 2255 | 2125 | 3848 | **2235** | 3867 | 2592 | 9957 |
| Interaction model | Explained deviance (%) | 42.4 | **58.9** | **51.5** | 24.3 | **33.8** | **47.3** | **35.6** |
| | PC 1 | *** | *** | *** | *** | *** | | *** |
| | PC 2 | * | | *** | * | | *** | *** |
| | PC | *** | *** | *** | ** | *** | ** | *** |
| | Distance to closest colony | - | *** | *** | *** | *** | *** | *** |
| | AIC | 2264 | **2115** | **3678** | 2237 | **3678** | **2512** | **9674** |
| AIC (global model) – AIC (interaction model) | | -9 | 10 | 170 | -2 | 189 | 80 | 283 |

### 1. Bottlenose dolphins

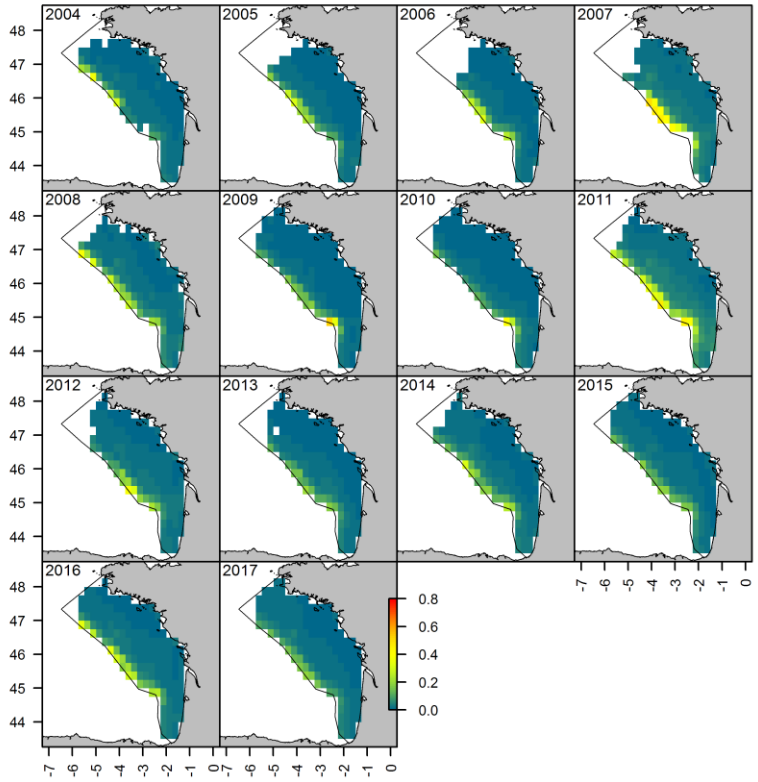


**Figure 3. Annual prediction for bottlenose dolphins, based on the global model.**

### 2. Black-legged kittiwakes

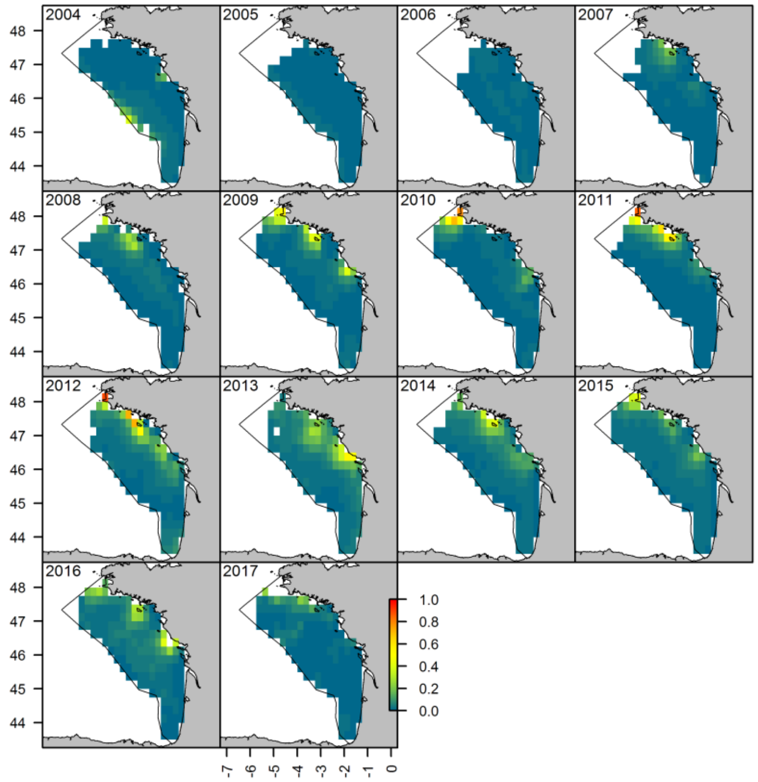


**Figure 4. Annual prediction for black-legged kittiwakes, based on the interaction model.**

### 3. Auks

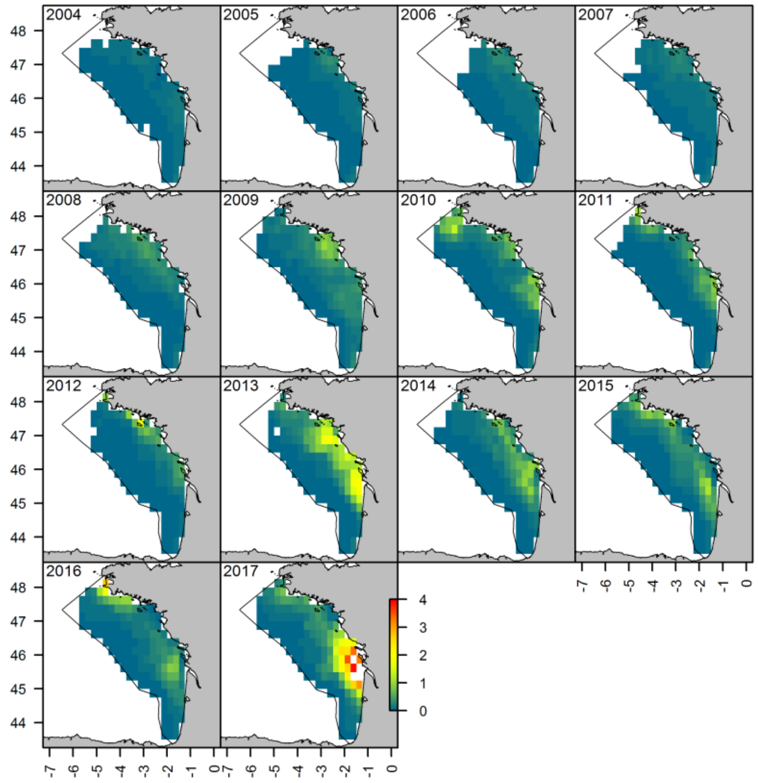


**Figure 5. Annual prediction for auks, based on the interaction model.**

### 4. Storm petrels

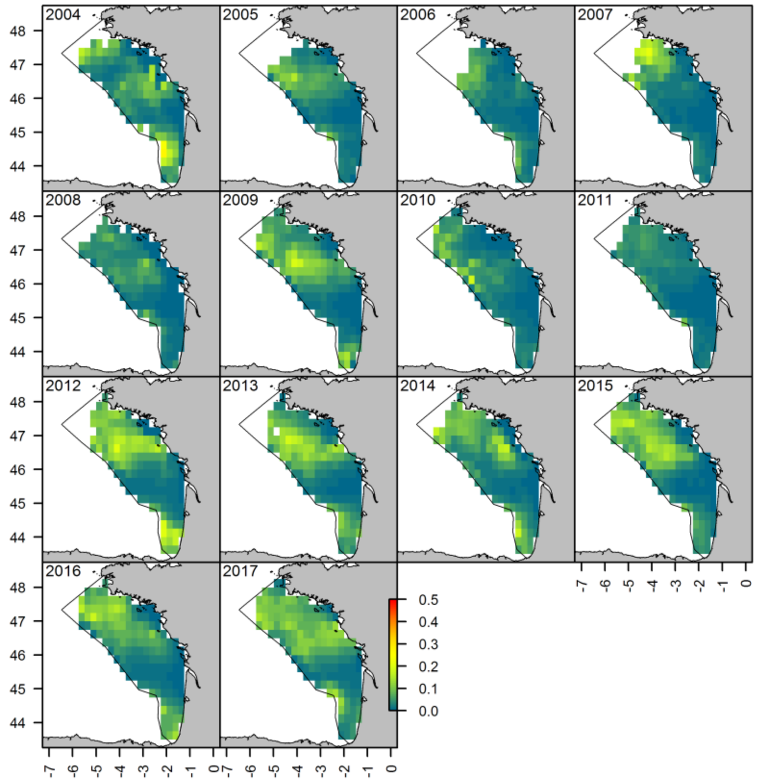


**Figure 6. Annual prediction for storm petrels, based on the global model.**

### 5. Northern fulmars

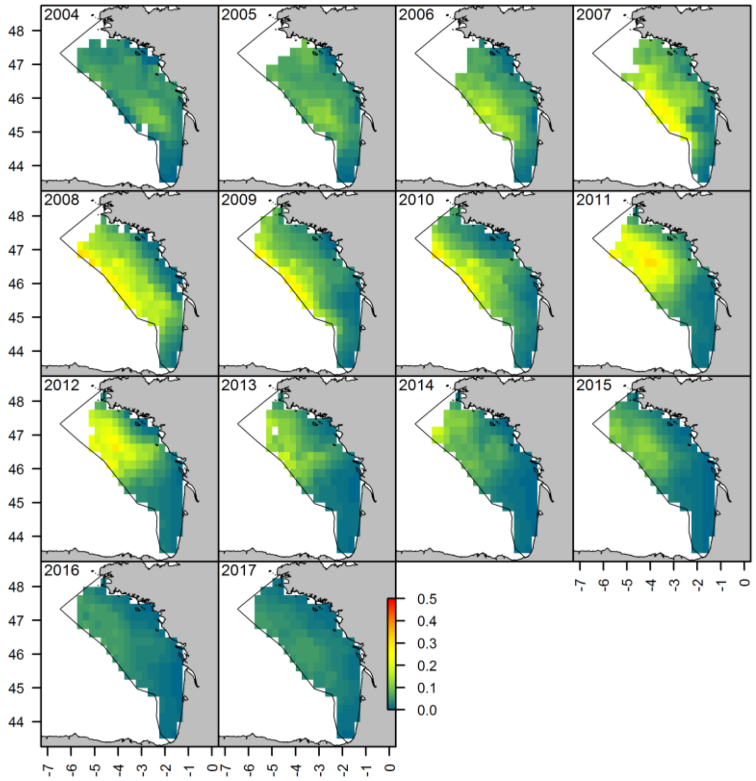


**Figure 7. Annual prediction for northern fulmars, based on the interaction model.**

### 6. Small-sized shearwaters

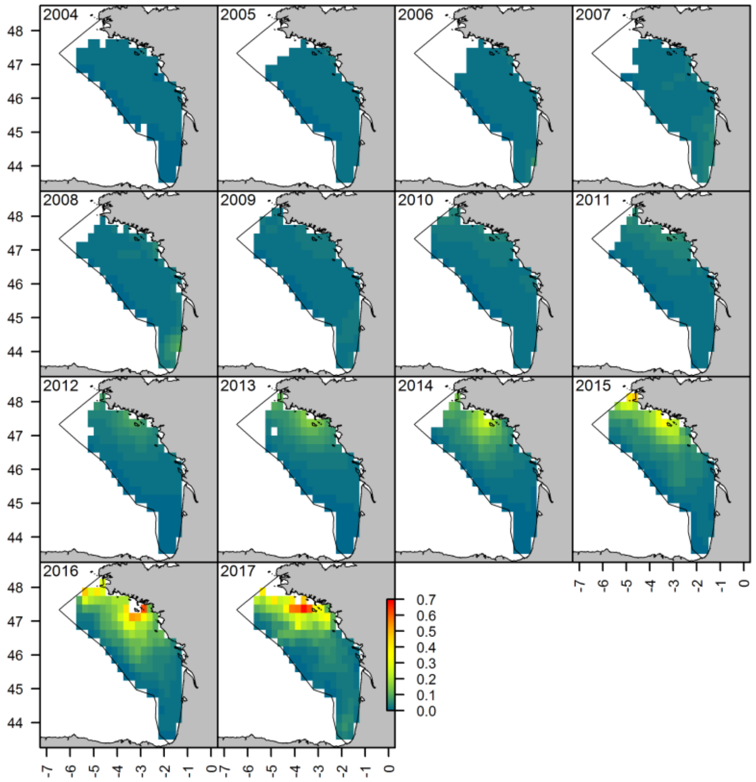


**Figure 8. Annual prediction for small-sized shearwaters, based on the interaction model.**

7. Northern gannets

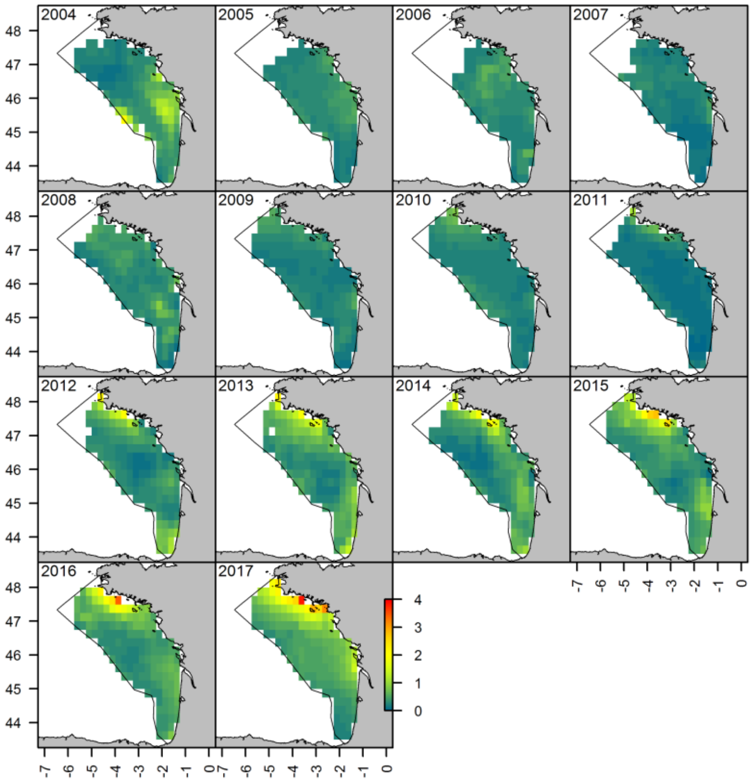


**Figure 9. Annual prediction for northern gannets, based on the interaction model**
