## Appendix B for "The persistence in time of distributional patterns in marine megafauna impacts zonal conservation strategies"

### Core areas of distribution

Based on predicted abundances, we identified the smallest sets of pixels including 50% of the population, highlighting core areas of distribution. Core areas are in black, the remaining study area with prediction in grey. See text for details.

1. <u>Bottlenose dolphin</u>

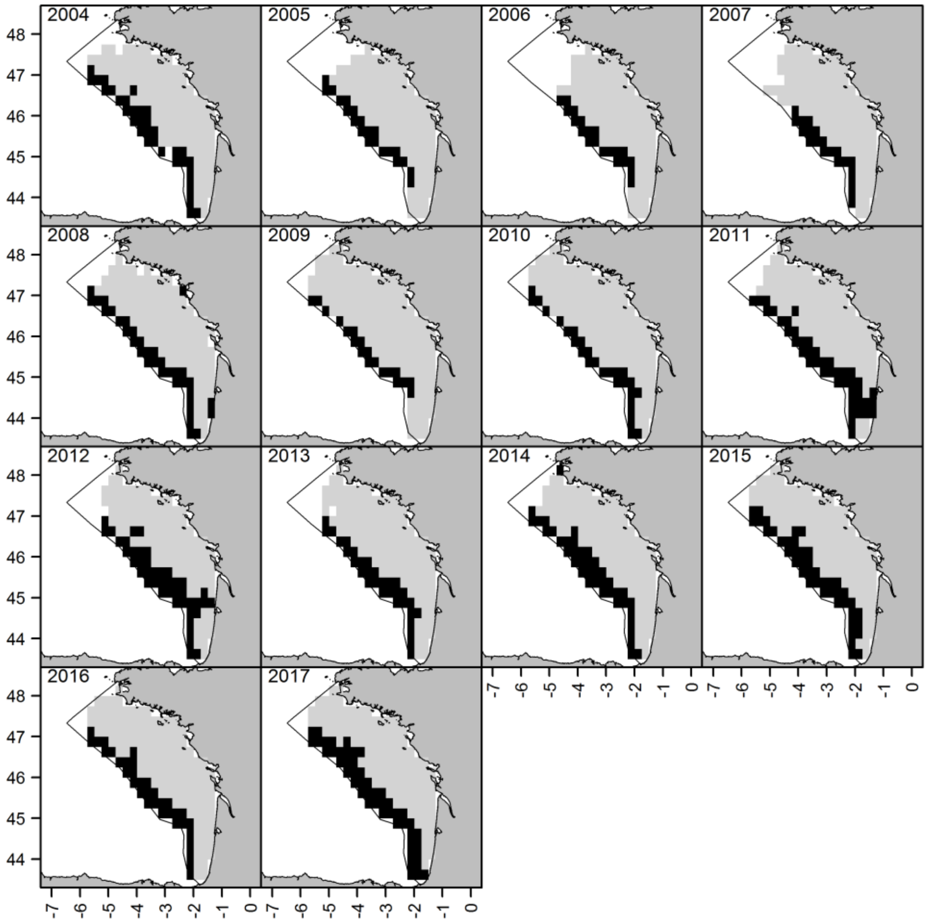

2. Black-legged kittiwake

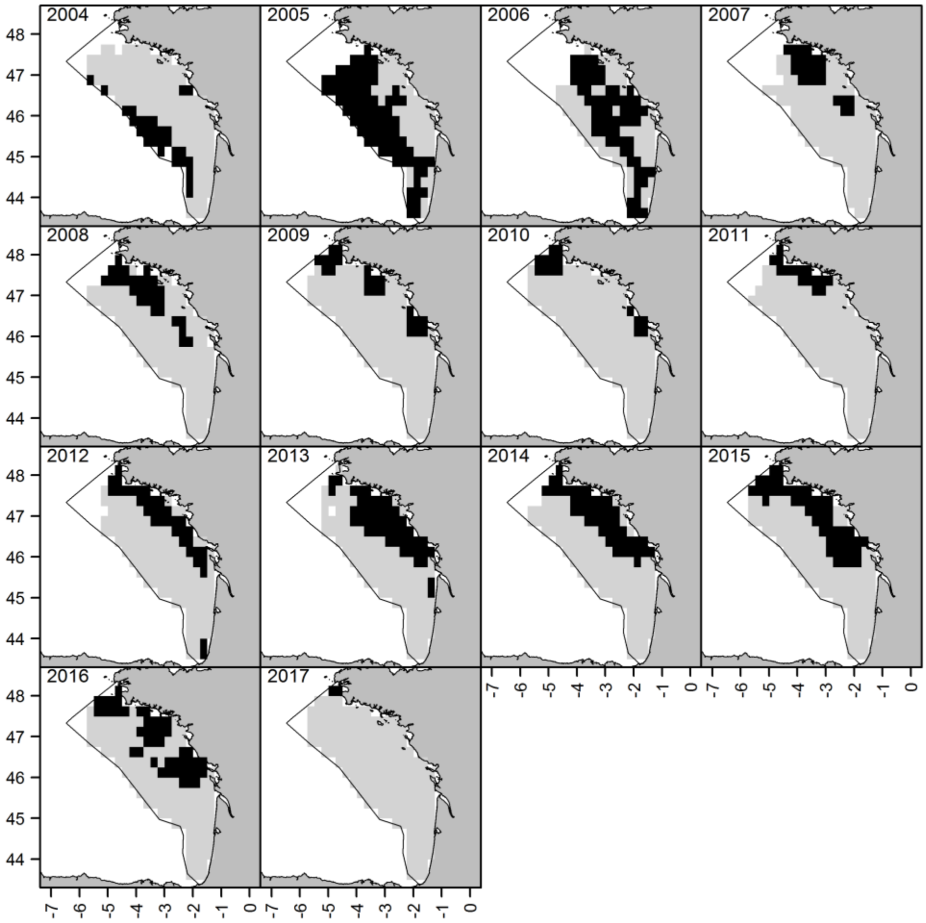


3. Auks

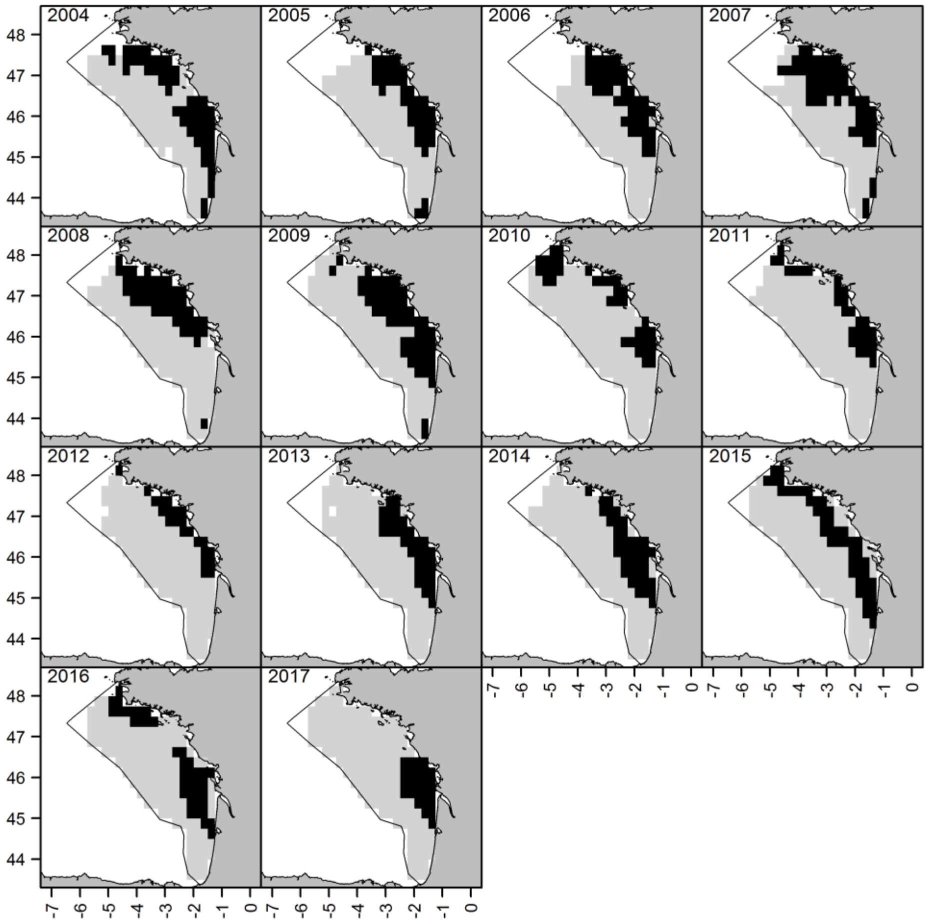

4. Storm petrels

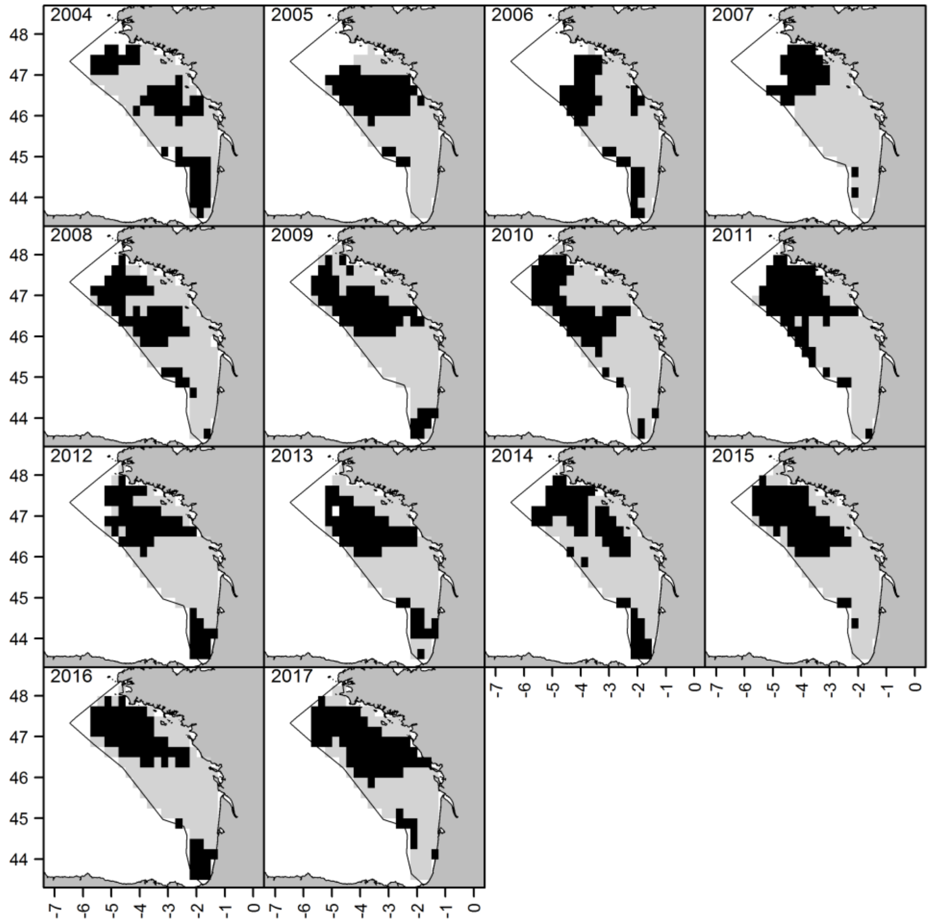


5. Northern fulmar

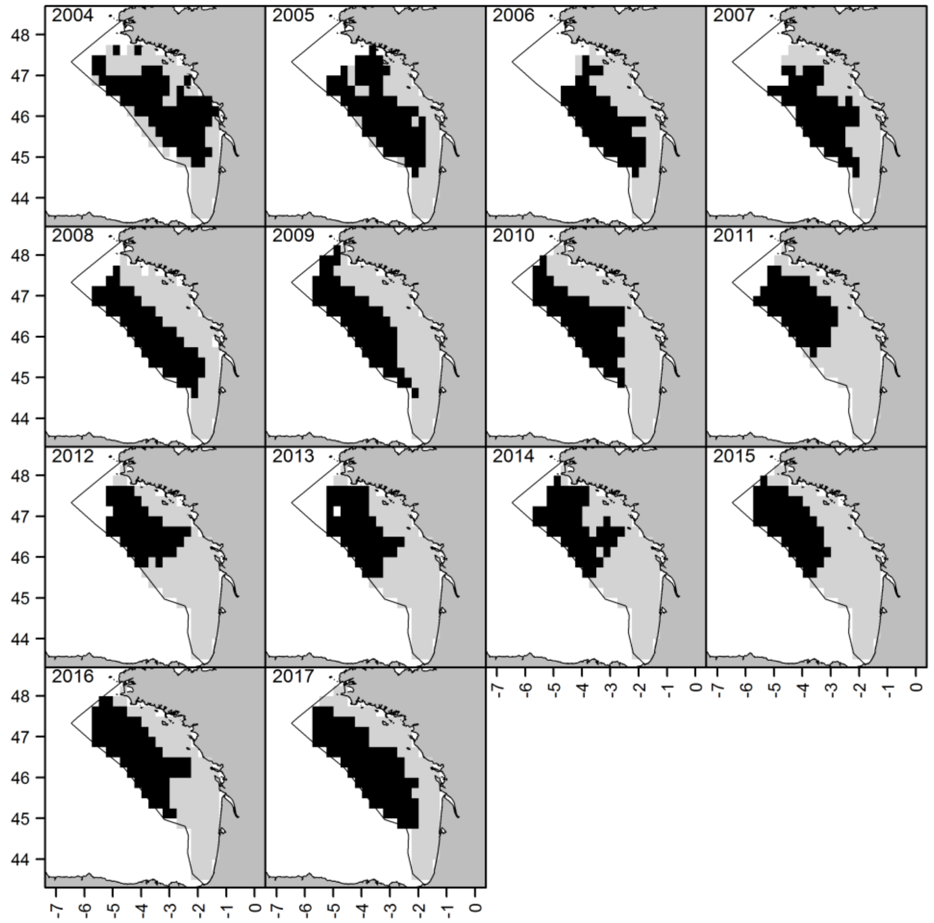

6. Small-sized shearwaters

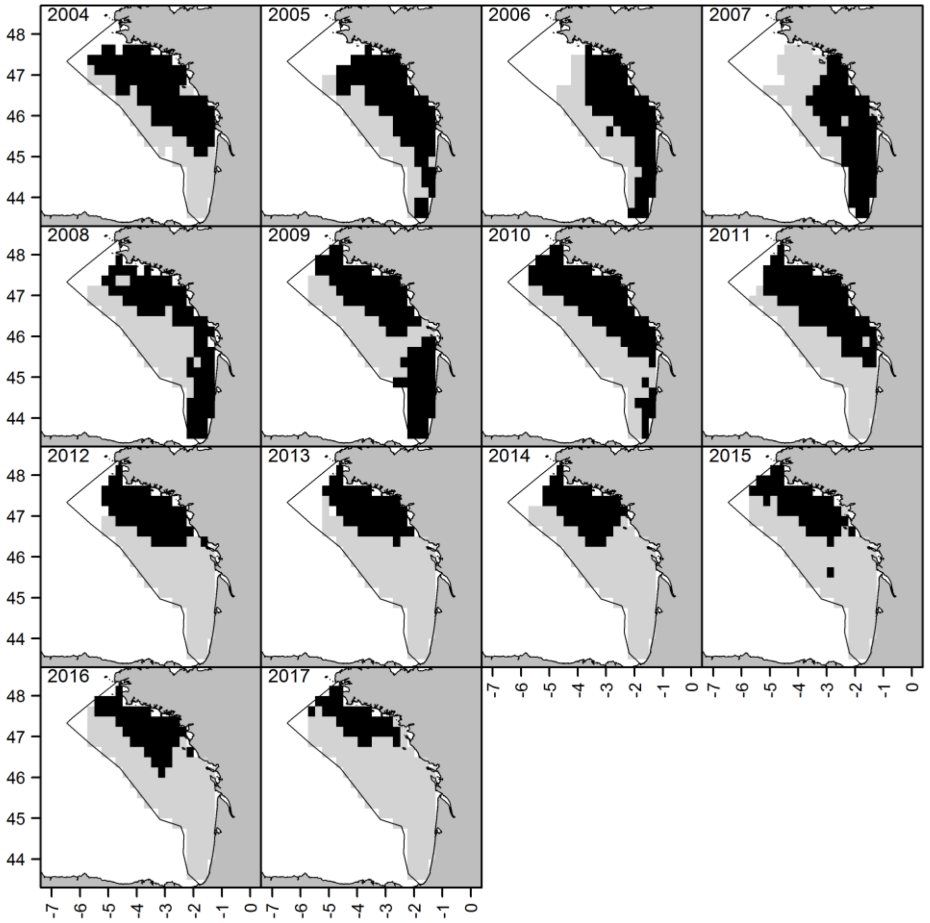


7. Northern gannet

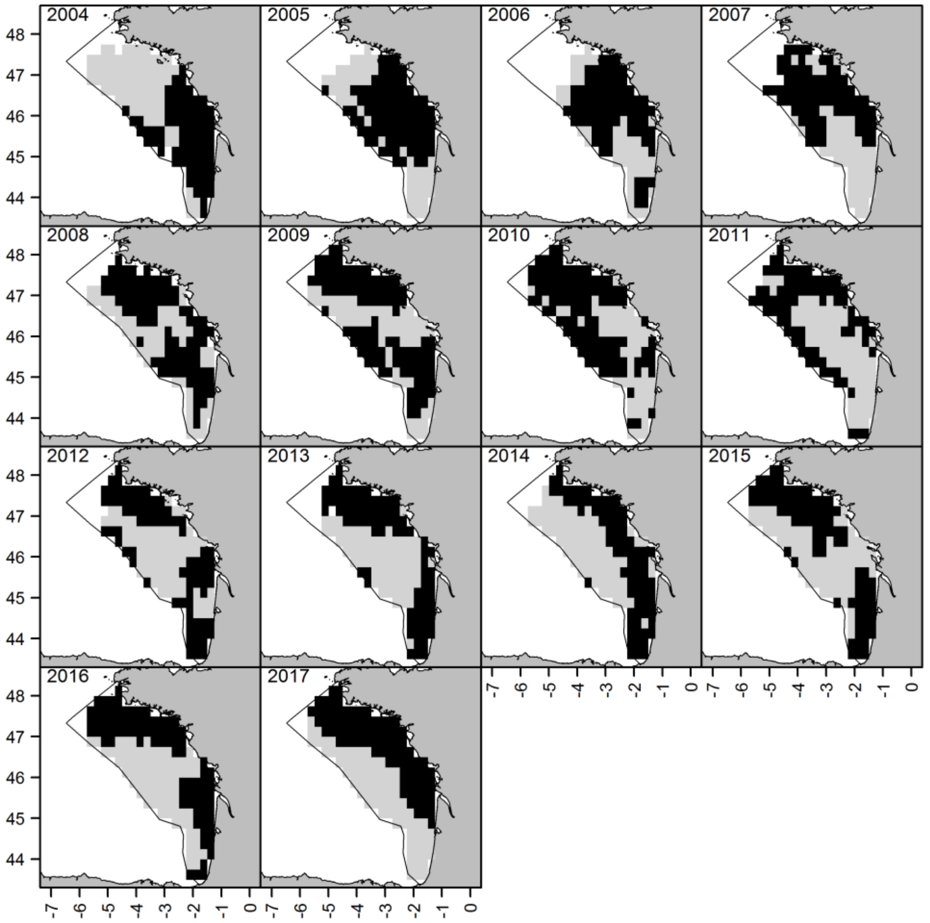
